# Single-molecule characterisation of soluble beta-amyloid aggregate binding by Aducanumab, Lecanemab, Gantenerumab, and Donanemab

**DOI:** 10.1101/2024.10.11.617910

**Authors:** Emre Fertan, Jeff Y. L. Lam, Giulia Albertini, Maarten Dewilde, Yunzhao Wu, Oluwatomi E. S. Akingbade, Dorothea Böken, Elizabeth A. English, Bart De Strooper, David Klenerman

## Abstract

Monoclonal antibodies Aducanumab, Lecanemab, Gantenerumab, and Donanemab have been developed for treatment of Alzheimer’s disease. Here, we have used single-molecule detection techniques and super-resolution imaging to characterise the binding of these antibodies to beta-amyloid aggregates including human post-mortem brain samples. Lecanemab is the best antibody in terms of binding to the small-soluble beta-amyloid aggregates, affinity, aggregate coating, and the ability to bind to post-translationally modified species, explaining its therapeutic success.

---

Alzheimer’s disease (AD) is a growing global pandemic^1^. The initiating role of beta-amyloid (Aβ) in AD pathology has led to the development of the amyloid-cascade hypothesis, postulating the accumulation and aggregation of Aβ40/42 peptides as the initial cause of AD^2,3^ and clearance of Aβ is a viable therapeutic strategy. The monoclonal antibodies (MAb) Aducanumab (Biogen), Lecanemab (BioArctic and Eisai), Donanemab (Eli Lilly), and Gantenerumab (Roche) targeting Aβ have been developed and recently tested in clinical trials.

Aducanumab is a recombinant human MAb that binds to amino acids 3-7 of the Aβ peptide^4^. Lecanemab is the humanised version of the murine mAb158 antibody, which primarily targets protofibrils^5^. Gantenerumab has two identified binding sites on the Aβ peptide; one closer to the N-terminal, targeting amino acids 3-11, and one later on in the middle of the peptide at the amino acids 18-27^6^. Donanemab targets the N-terminal truncated pyroglutamate Aβ peptide at position 3 (pGlu3-Aβ)^7^.

While Aducanumab, Lecanemab, and Gantenerumab all primarily target small-soluble aggregates of Aβ, of these three MAbs Lecanemab has shown the most desirable outcomes in clinical trials in terms of Aβ clearance and cognitive performance^8^, yet the reason of this has not been fully understood. All three MAbs have a preference for aggregated Aβ over monomeric species, and while Aducanumab and Gantenerumab show stronger binding to fibrillar aggregates, Lecanemab shows the strongest binding to protofibrils of Aβ^9^. Meanwhile, Donanemab has also been shown to slow-down the progression of AD significantly at early symptomatic AD cases with comparable success to Lecanemab^10^, suggesting that targeting pGlu3-Aβ may also be a worthwhile strategy. The small-soluble aggregates primarily targeted by these MAbs are below the diffraction-limit of light, highly heterogenous in terms of morphology, and exist in low concentrations, making them challenging to study, so our group has developed single-molecule detection techniques to characterise them^11–13^. Here, we used single-molecule array (SiMoA) and super-resolution microscopy (**Figure 1**) to investigate the Aβ and pGlu3-Aβ aggregate binding properties of Aducanumab, Lecanemab, Gantenerumab, and Donanemab, and provide explanations for the therapeutic superiority of Lecanemab and Donanemab.

**Figure 1:**
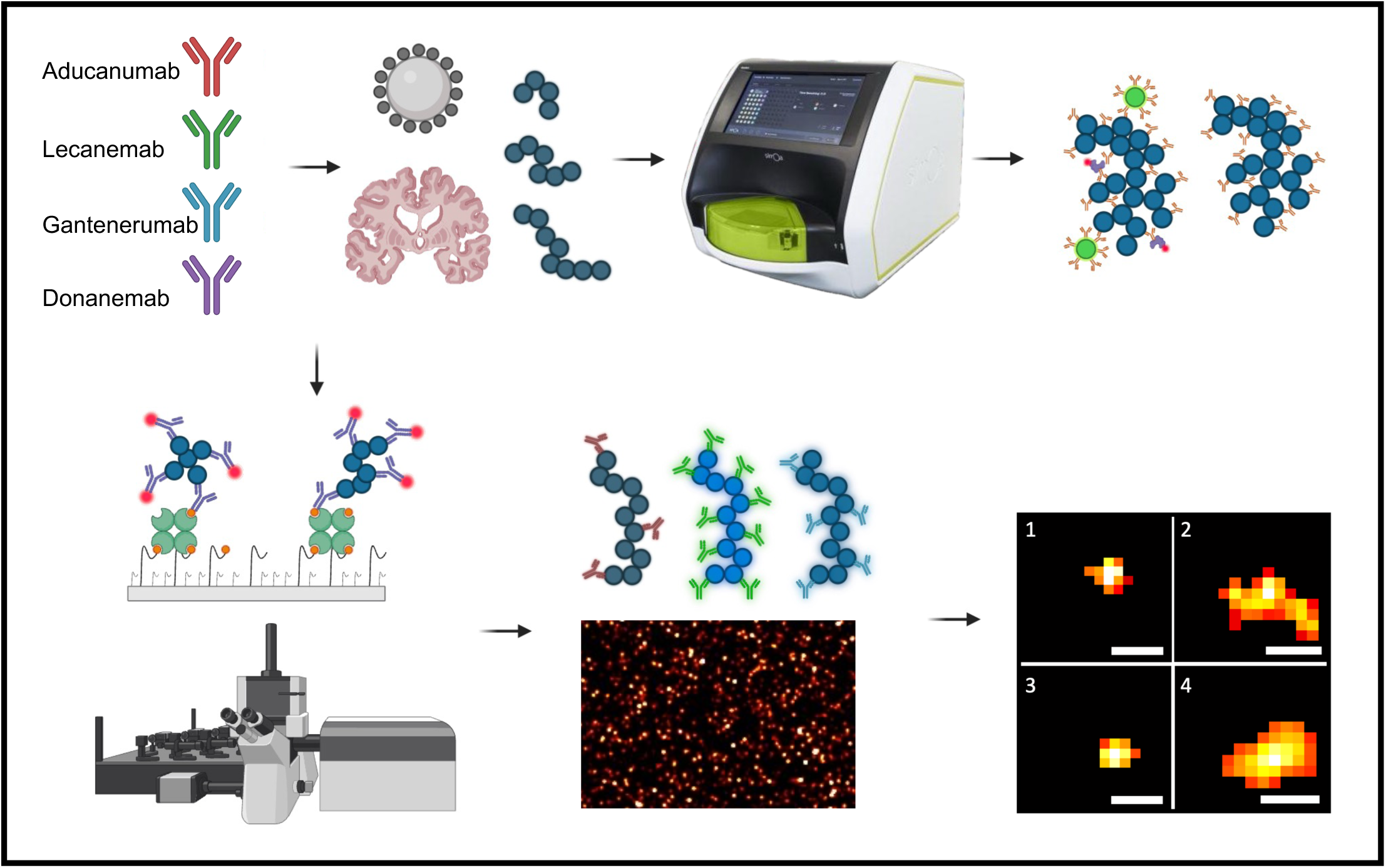
Visual summary of the experiments. Monoclonal antibodies were produced in-house from publicly available sequences and tested with silica nanoparticles coated with beta-amyloid, *in-vitro* aggregates, and human post-mortem Alzheimer’s disease brain homogenates using SiMoA, single-molecule pulldown, and *d*STORM. The SiMoA platform utilises paramagnetic beads coated with capture antibodies, binding to targets of interest at ultra-low concentrations, forming a small number of immunocomplexes on the beads, which are then detected using streptavidin beta-galactosidase and resorufin beta-D-galactopyranoside interactions, providing a binary (digital) readout for aggregate quantification. Direct stochastic optical reconstruction microscopy (*d*STORM) along with single-molecule pulldown (SiMPull) uses a glass surface optimised to capture targets of interest with high sensitivity, which are then imaged with a resolution limit of 30 nm (under the diffraction limit of light), allowing the morphological characterisation of soluble aggregates (scale bars are 50 nm).

We started our investigations by comparing the signal from the MAbs using 15-nm silica nanoparticles conjugated to monomeric Aβ42 or pGlu3-Aβ, mimicking aggregates with a known size and shape (**Figures 2A-B**). When tested with the Aβ42 coated beads, the signal correlated positively with bead concentration for all the MAbs except for Donanemab, which showed a consistently low signal confirming that Donanemab does not bind to full-length Aβ42 peptide (**Figure 2A**). In the samples without any coated nanoparticles (i.e. blank controls), the highest background signal was observed from Gantenerumab and the lowest from Lecanemab. Lecanemab also showed the steepest rise in signal with increasing concentrations of nanoparticles. The limit of detection (LoD) of Aducanumab, Lecanemab, and Gantenerumab were 200, 2, and 70 picomolar (pM), respectively. When we repeated this experiment with pGlu3-Aβ coated silica nanoparticles, Donanemab did show a signal as expected, which increased with the sample concentration, showing that it binds specifically to pyroglutamate-modified species (**Figure 2B**). Interestingly, Lecanemab and (to a much lower extent) Aducanumab also showed a signal with the pGlu3-Aβ coated silica nanoparticles, showing that these MAbs also bind to pGlu3-Aβ, unlike Gantenerumab. The LoD of Aducanumab, Lecanemab, Gantenerumab, and Donanemab for pGlu3-Aβ coated silica nanoparticles were 56, 4, 220, and 5 pM, respectively.

**Figure 2:**
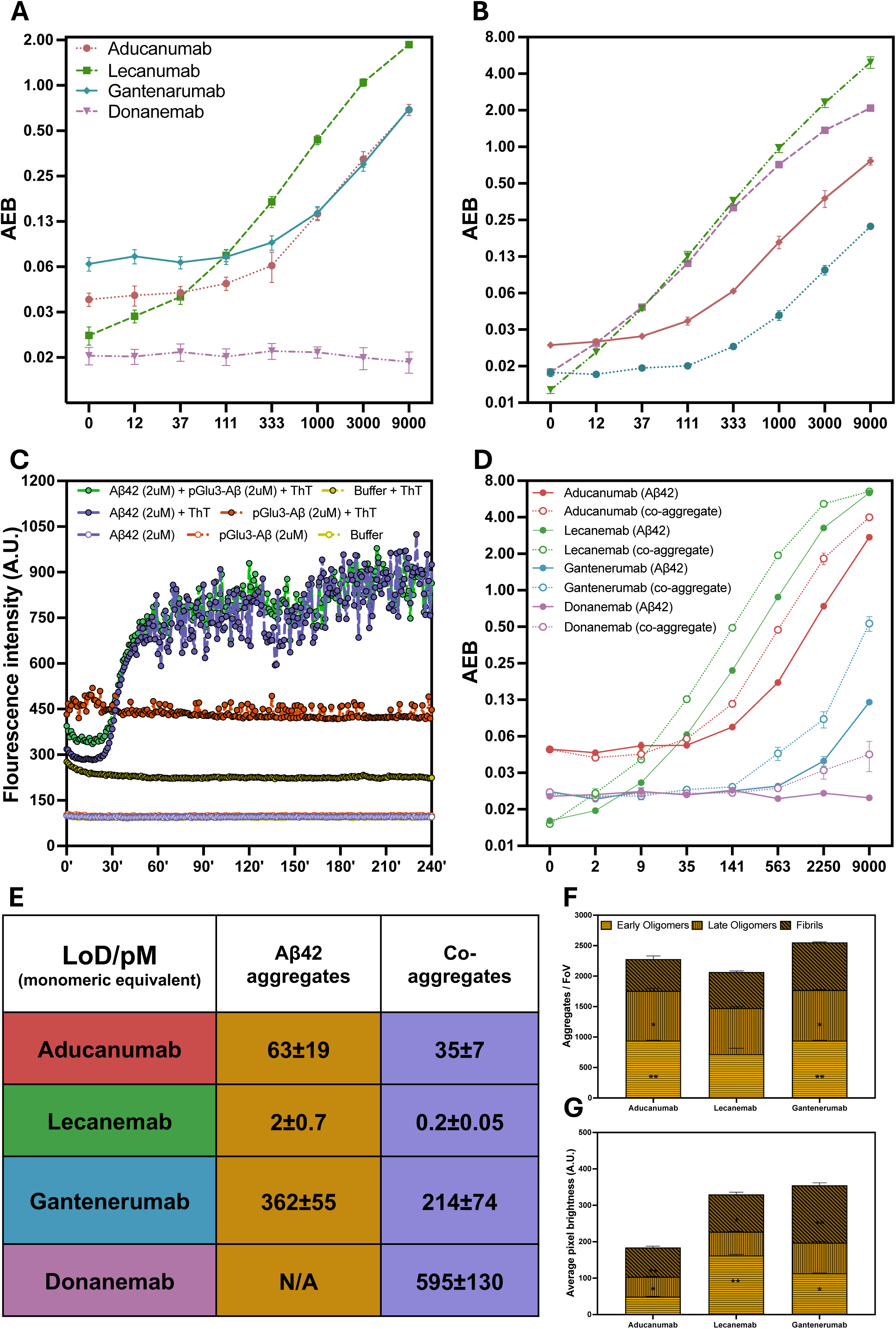
Characterisation of the monoclonal antibodies using synthetic aggregates. Signal on SiMoA was compared between Aducanumab, Lecanemab, Gantenerumab, and Donanemab using beta amyloid 42 (**A**) and pyroglutamate beta-amyloid (**B**) coated glass nanoparticles at different concentrations (x-axis shows concentration in nM). Each antibody-bead concentration combination was tested at least on four independent wells. For beta-amyloid 42 coated beads, a significant antibody by concentration interaction was determined (AIC = 470.4, F = 306.6, p < 0.001, ηp^2^ = 0.855). While Donanemab did not show a dose dependent increase in signal, Aducanumab, Lecanemab, and Gantenerumab all showed a significant increase with increased bead concentration, with the greatest increase seen in Lecanemab (**A**). For pyroglutamate beta-amyloid coated beads, once again a significant antibody by concentration interaction was present (AIC = 101.3, F = 264.4, p < 0.001, ηp^2^ = 0.860). All four antibodies showed a concentration-dependent increase in signal, with the greatest signal acquired from Lecanemab and Donanemab, while the weakest signal was measured from Gantenerumab (**B**). In addition to coated silica-nanoparticles, *in-vitro* aggregates of beta-amyloid were also prepared by incubating beta-amyloid 42 and pyroglutamate beta-amyloid for different durations. A thioflavin T assay was performed to characterise the aggregation dynamics of different species, revealing the formation of thioflavin T-positive aggregates over time in the samples containing pure beta-amyloid 42 and a mixture of beta-amyloid 42 and pyroglutamate beta-amyloid, however the signal did not change for the pure pyroglutamate beta-amyloid signal, showing a lack of aggregation (**C**; x-axis shows time in minutes). Then these samples were tested on SiMoA on three independent wells, revealing a three-way interaction between the antibody, aggregate type, and concentration (AIC = 584.2, F = 23.7, p < 0.001, ηp^2^ = 0.797). Overall, the signal with the mixed aggregates was higher for all antibodies than the pure beta-amyloid aggregates and the signal increased with aggregate concentration. The highest signal was detected by Lecanemab, followed by Aducanumab, Gantenerumab, and Donanemab (**D**; x-axis shows concentration in nM). The limit of detection was in parallel to the signal at high-concentrations (**E**). The performance of Aducanumab, Lecanemab, and Gantenerumab with *in-vitro* aggregates at different sizes were also compared using single-molecule pulldown (**F-G**), with imaging the aggregates on two independent wells for at least nine fields of view per well. For the number of aggregates detected, there was an antibody by aggregate type interaction (AIC = 800.0, F = 4.9, p = 0.002, ηp^2^ = 0.214). The greatest number of aggregates were detected with Gantenerumab, followed by Aducanumab, and Lecanemab. While Aducanumab and Lecanemab showed a strong preference for early and late oligomers, the difference was smaller for Gantenerumab (**F**). Aggregate brightness was also measured, with controlling for the number of dyes per antibody, providing a measure for number of antibodies binding to unit area of each aggregate (i.e. the aggregate coating ability). Again, an antibody by aggregate type interaction (AIC = 703317.0, F = 91.8, p < 0.001, ηp^2^ = 0.006) was found, Lecanemab achieved the highest brightness of all antibodies when tested with early oligomers, which was significantly higher than the signal it provided with fibrils and least with late oligomers. On the other hand, Gantenerumab and Aducanumab provided the brightest signal with fibrils, with the signal from Aducanumab being significantly dimmer than Gantenerumab, collectively showing a strong preference for early oligomers by Lecanemab, and a preference for fibrillar aggregates by Gantenerumab, and to a lesser extent Aducanumab (**G**).

Then, we aggregated synthetic monomeric Aβ at different time-points to compare the binding dynamics of the MAbs to different types of aggregates. First, we determined the aggregation rates of Aβ42 and pGlu3-Aβ with a thioflavin T (ThT) assay (**Figure 2C**). Starting with monomeric Aβ42, pGlu3-Aβ, or their mixture at 2 μM concentration, we observed that the ThT signal does not change for pure pGlu3-Aβ but increases for Aβ42 and the mixed sample, showing that pGlu3-Aβ does not aggregate on its own. We aggregated Aβ42 at 37℃ for 30-, 60-, and 180-minutes to produce “early oligomers”, “late oligomers”, and “fibrils”^14^. Since Donanemab does not bind to full length Aβ42 aggregates and pGlu3-Aβ does not aggregate on its own, we also produced co-aggregates of Aβ42 and pGlu3-Aβ aggregating a mixture of 2 µM Aβ42 and pGlu3-Aβ for 60-minutes (the late oligomer time-point) to compare Donanemab with the other MAbs. The size of these aggregates were determined using *d*STORM with the 6E10 antibody, confirming that the early oligomers were shortest (average length 105 nm) followed by the late oligomers (average length 113 nm) and the longest aggregates were the fibrils (average length 123 nm), while the co-aggregates of Aβ42 and pGlu3-Aβ were slightly larger.

Aducanumab, Lecanemab, and Gantenerumab showed a concentration-dependent signal with the late Aβ42 oligomers, while Donanemab did not show a signal above background, again confirming no binding to full-length Aβ42 (**Figure 2D**). Meanwhile, the signal was higher for the co-aggregates compared to the pure Aβ42 aggregates for Aducanumab, Lecanemab, and Gantenerumab. Given that Gantenerumab did not bind to pGlu3-Aβ coated silica-nanoparticles, this increase in signal may not be due to the presence of available pGlu3-Aβ epitopes, but due to the formation of larger aggregates or conformational changes taking place during co-aggregation, which increase epitope availability. Surprisingly, Donanemab did not give a significant signal with the co-aggregates, suggesting that the pGlu3-Aβ is not available for MAbs to interact with in the co-aggregates, potentially forming the core of the co-aggregate. Both with the pure Aβ42 and mixed Aβ42/pGlu3-Aβ aggregates, the lowest LoD was achieved by Lecanemab, followed by Aducanumab and Gantenerumab (**Figure 2E**).

We further tested the pure Aβ aggregates at different time points (early oligomers, late oligomers, fibrils) to identify the binding preferences of Aducanumab, Lecanemab, and Gantenerumab on single-molecule pulldown (SiMPull), comparing the number of aggregates detected by each MAb, since Donanemab showed no binding. Regardless of aggregate type, the greatest number of aggregates were detected by Gantenerumab, followed by Aducanumab, and Lecanemab (**Figure 2F**). While Aducanumab and Gantenerumab showed a preference for early oligomers, followed by late oligomers, and fibrils, Lecanemab did not show a size-dependent aggregate specificity and detected similar number of aggregates in all the samples. However, when we compared the average pixel brightness (normalised to the labelling efficiency), which is a function of number of antibodies binding to an aggregate (i.e. aggregate coating ability) the brightest aggregates bound by Aducanumab and Gantenerumab were the fibrils, while Lecanemab had the brightest signal with the early oligomers (**Figure 2G**). The aggregates detected by Lecanemab and Gantenerumab were significantly brighter than those detected by Aducanumab.

We then tested the MAbs on soluble Aβ aggregates harvested from homogenised human middle-temporal gyrus (Brodmann’s area 21) samples with Braak stage 0, 3, and 5 (B0, B3, B5) pathology. In the SiMoA analysis, Aducanumab and Gantenerumab showed the largest signal with the B5 samples (**Figure 3A**), suggesting the presence of aggregates to which they preferentially bind. The largest signal with Lecanemab was detected with the B3 samples, showing a preference for aggregates formed during the early disease stages. Remarkably, Donanemab showed a consistently low signal with samples from all stages, indicating a low epitope availability of the pGlu3-Aβ *in-vivo* as well. These results suggest that the reason of Donanemab’s therapeutic success is not through binding to the small-soluble aggregates in the AD brain.

**Figure 3:**
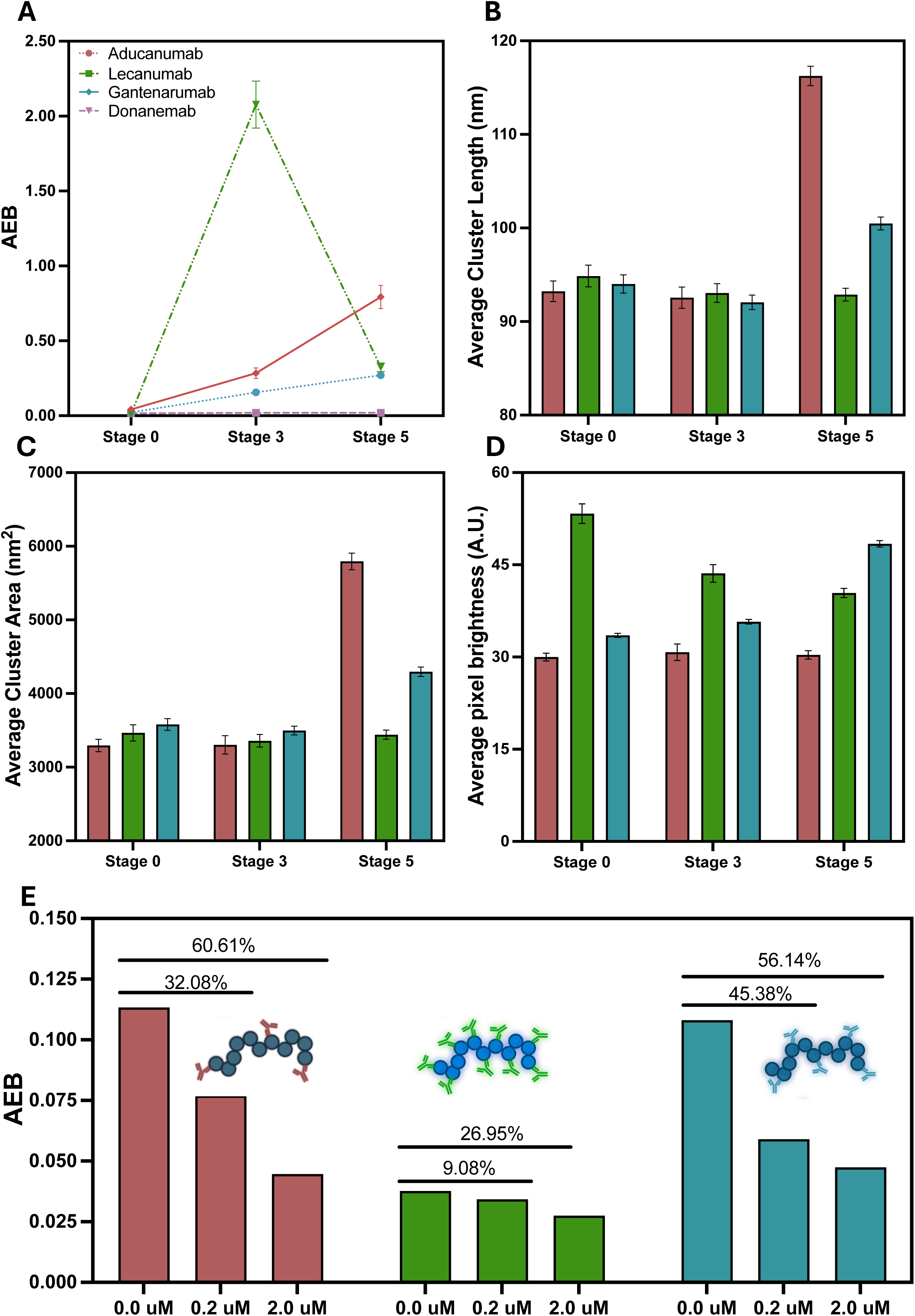
Characterisation of the monoclonal antibodies using post-mortem human Alzheimer’s disease brain homogenates. Signal on SiMoA was compared between Aducanumab, Lecanemab, Gantenerumab, and Donanemab using human post-mortem brain homogenates from patients with Braak stage 0, 3, or 5 Alzheimer’s disease, with two independent brain samples tested in two independent wells per condition (**A**). A significant antibody by Braak stage interaction was observed (AIC = 63.19, F = 128.1, p < 0.001, ηp^2^ = 0.964); while Aducanumab and Gantenerumab signal positively correlated with the Braak stage, Lecanemab showed the highest detection with the Stage 3 samples. Meanwhile, Donanemab signal was at the baseline for the brain homogenates containing soluble beta-amyloid aggregates. Brain homogenates were also tested on single-molecule pulldown with *d*STORM, to measure the length (**B**) and area (**C**) of the aggregates detected by the antibodies on two independent wells with at least two fields of view each. There was an antibody by Braak stage interaction for both aggregate length (AIC = 231669.0, F = 54.9, p < 0.001, ηp^2^ = 0.008) and area (AIC = 497262.0, F = 61.7, p < 0.001, ηp^2^ = 0.008). All antibodies showed a size preference and mostly bound to the aggregates 90 nanometres in length, yet the average length of the detected aggregates increased at Braak stage 5 for Aducanumab and Gantenerumab, while Lecanemab remained preferentially binding to the smaller aggregates at this stage. Average aggregate brightness as also measured during diffraction-limited imaging (same brain samples imaged in two independent wells with nine fields of view), with controlling for the number of dyes per antibody, providing a measure for number of antibodies binding to unit area of each aggregate (**D**). While the brightness signal was provided by Lecanemab overall and on average the Stage 5 brains were brighter, Lecanemab actually provided the brightest signal with the Stage 0 samples, followed by Stage 3, showing a preference for aggregates formed at early disease stages, unlike Gantenerumab and Aducanumab, which had a brighter signal with the Stage 5 samples, resulting in an antibody by Braak stage interaction (AIC = 466760.0, F = 98.6, p < 0.001, ηp^2^ = 0.007). Lastly, an immuno-pulldown was performed using the antibodies to identify the amount of available binding regions on the aggregates they detect (**E**). The greatest signal reduction was seen in Aducanumab, followed by Gantenerumab, and Lecanemab, once again showing the superior aggregate coating ability of Lecanemab.

We then performed *d*STORM imaging on these samples to measure the size of the aggregates which Aducanumab, Lecanemab, and Gantenerumab binds to. While the binding preference of the MAbs towards brain-derived aggregates at different sizes did not differ significantly at the earlier stages when the aggregates are smaller, Aducanumab had a greater affinity towards larger aggregates formed at later stage AD (**Figure 3B and 3C**). Once again using average aggregate brightness, we validated the epitope availability and the efficiency of the MAbs at coating the aggregates at different Braak stages. Regardless of MAb, the brightest aggregates were observed in the B5 samples, confirming that the larger aggregates at this stage have a greater number of binding sites (**Figure 3D**). Lecanemab showed the brightest signal overall, and the signal intensity was Braak stage dependent: for the B0 and B3 samples, Lecanemab provided the highest signal intensity. However, for the B5 samples, Gantenerumab had the brightest signal, once again suggesting that Lecanemab preferentially binds the smaller aggregates formed during earlier AD stages and more epitopes for Gantenerumab become available as the aggregates grow larger.

To further test the binding affinity of the MAbs to brain-derived soluble Aβ aggregates and their capacity to coat these aggregates, we performed an immuno-blocking experiment using SiMoA. Samples were first incubated with the MAbs at low (11.8) or high (118) micromolar (µM) concentration and then the SiMoA assay was performed using the same MAb for capture, with the rationale that if the aggregate has more binding site for a specific MAb, then the antibodies on the bead used in the SiMoA assay will have high avidity and be able to capture aggregates even if there is a high concentration of MAb in solution (**Figure 3E**). Agreeing with the brightness data, the least amount of change was observed in the signal from Lecanemab, and a much greater reduction in signal was observed with Aducanumab and Gantenerumab. Collectively, these results are showing that Lecanemab has more binding regions than Gantenerumab and Aducanumab on the soluble Aβ aggregates, especially during the earlier disease stages when the soluble Aβ aggregates are smaller.

Clinical studies have shown that monoclonal antibodies are a useful tool to clear Aβ from the brains of AD patients, which does not only lead to reduced plaque load, but also reduced pathological tau accumulation and improvements in cognitive decline^10,15^. However, the possible mechanism(s) of action of these MAbs have not been fully understood. Aducanumab has not met the desired efficiency^16^ and will be discontinued within 2024. Meanwhile, Gantenerumab failed to achieve significant therapeutic success in clinical trials^17^. On the other hand, Lecanemab and Donanemab were both approved by the FDA following desirable clinical outcomes of significant reduction in amyloid-PET signal and cognitive improvement. While our data fails to explain the success of Donanemab in Aβ clearance, it provides significant insights for the success of Lecanemab.

The main differences between Lecanemab and the other MAbs is that it (1) preferentially binds smaller Aβ aggregates, (2) with more antibodies per aggregate, and (3) it has a significantly higher affinity for aggregates by an order of magnitude than Aducanumab and Gantenerumab. Given the antibody concentration in the brain is likely to be low^18^, a higher affinity would lead to more binding and removal of smaller aggregates. Plaque dissociation may then occur due to the shift in the dynamic equilibrium between aggregates and plaques, although other mechanisms cannot be excluded. However, the comparable clinical benefits of Donanemab cannot be explained by soluble aggregate binding, so other mechanism such as binding to the insoluble plaques or triggering microglial phagocytosis must therefore be important. It would be valuable to characterise these in future studies. Lastly, we also showed that Lecanemab and Aducanumab show binding to the pGlu3-Aβ, which has not been previously reported and may contribute to their therapeutic success.

## Declarations

### Ethical approval

The post-mortem human brain samples were used with the approval of the London—Bloomsbury Research Ethics Committee (16/LO/0508) and the Edinburgh Brain Bank, which has ethical approval from the East of Scotland Research Ethics Service to function as a research tissue bank (REC 21/ES/0087). The brain samples were voluntarily donated without any compensation. We would like to thank the donors and their families.

### Availability of data and materials

Data collected during these experiments will be made available upon reasonable request from the corresponding author.

### Author contributions

**J.Y.L.L.**: Conception and design, data collection and analysis, manuscript preparation. **E.F.**: Conception and design, data collection and analysis, data analysis, manuscript writing. **G.A.**: Conception and design, preparation of monoclonal antibodies, and manuscript preparation. **M.D.**: Preparation of the monoclonal antibodies. **Y.W., O.E.S.K.**: Data collection. **D.B., E.A.E.**: Sample preparation. **B.D.S.**: Conception and design, manuscript preparation, supervision of the project. **D.K.**: Conception and design, data interpretation, manuscript writing, supervision of the project.

## Online methods

### Preparation of Aducanumab, Lecanemab, Gantenerumab and Donanemab

The amino acid sequences for the variable domains of Aducanumab, Lecanemab, Gantenerumab and Donanemab were retrieved from the KEGG DRUG Database. Synthetic genes encoding for the respective variable domains, preceded by the mouse Ig heave leader signal, were ordered at Twist Biosciences (South San Francisco, CA, USA), and cloned in pTRIOZ-hIgG (Invivogen, San Diego, CA, USA). The VL domain was cloned by using restriction enzymes AscI/BsiWI, the VH domain by AgeI/NheI. The antibody encoding open reading frames were sequence confirmed by Sanger sequencing (LGC Genomics, Berlin, Germany). Plasmid DNA was delivered to CHO hamster ovarian cells (A29127, Thermofisher) by transient transfection with the TransIT-PRO® Transfection Reagent (MIR 5740, Mirus Bio) according to manufacturer’s protocol. Transfected CHO cells were cultured for 14 days in suspension on agitation at 32°C. Two weeks post-transfection, cell supernatants were collected and incubated overnight at 4°C with AmMag^TM^ Protein A Magnetic Beads (L00939, GenScript). The beads were collected using a magnetic separation rack and targeted antibodies were separated using the AmMag™ SA Plus system (L01013, GenScript). The purity of Aducanumab, Lecanemab, Gantenerumab and Donanemab was estimated to be above 75% by densitometric analysis of the Coomassie Blue-stained SDS-PAGE gel under non-reducing conditions.

### Preparation of *in-vitro* aggregates

Lyophilised monomeric Aβ42 peptide (Abcam, Cat. No. ab120301, Lot No. APN22027-1-1) and pGlu3-Aβ peptide (Anaspec, Cat. No. AS-29907-01, Lot No. 2355657) were dissolved in PBS (ThermoFisher Scientific, Cat. No. 10010023) at 20 μM on ice. The solutions were quickly aliquoted and snap frozen. To prepare *in-vitro* Aβ42 and co-aggregates (mixed Aβ42 and pGlu3-Aβ), an aliquot was thawed on ice and diluted to have 2 μM final concentration of each peptide, in PBS. The diluted solution was then aliquoted (100 μL/aliquot) and incubated at 37 °C under quiescent conditions for 30 min, 60 min and 180 min as early oligomers, late oligomers, and fibrils respectively. The aggregation process was monitored using 20 μM (as confirmed by A_412_ (ε_412_ = 31,600 M^−1^cm^−1^)) ultrapure-grade Thioflavin-T (ThT, AnaSpec, Cat. No. AS-88306; **Supplemental Figure 1A**). Thereafter, the *in-vitro* aggregates were aliquoted (50 μL) and snap frozen. The aliquots were stored at −80°C until use.

### Preparation of Aβ aggregate-mimics

The synthetic Aβ42 and pGlu3-Aβ aggregate-mimics were prepared using a published protocol^1^ with modifications. To 500 µL of carboxylated silica nanoparticles (cSiNaP) (particle size with 15-nm diameter, 11.47 µM in dimethylformamide (DMF), Merck, Cat. No. 660450), 500 µL of 18.2-MΩ cm water was introduced. The mixture was then centrifuged (10,000 g, room temperature, 1 h) and the pellet was resuspended with 500 µL of MES buffer (2-(N-morpholino)ethanesulfonic acid) buffer, 10 mM, pH 5.7). Meanwhile, 1-ethyl-3-(3-dimethylaminopropyl) carbodiimide (EDC, ThermoFisher, Cat. No. A35391) and sulfo-N-hydroxysuccinimide (sulfo-NHS, ThermoFisher, Cat. No. A39269) were freshly dissolved in cold MES buffer (10 mM, pH 5.7) at 52.16 mM (10 mg/mL) and 92.11 mM (20 mg/mL), respectively. The cSiNaP was then diluted to 100 nM in MES buffer. EDC and sulfo-NHS were then introduced into the diluted cSiNaP suspension such that the mixture contained 400 µM EDC and 100 µM sulfo-NHS. The reaction mixture was sonicated for 30 min and then centrifuged (10,000 g, room temperature, 1 h). The pellet, i.e., activated SiNaP, was resuspended in fresh 10 mM MES buffer to give 200 nM suspension. To 1 mL of the activated SiNaP suspension, 1 mL of monomeric Aβ42 peptide or pGlu3-Aβ peptide solution (20 μM in PBS) was introduced. The reaction mixture was placed on a revolver rotator for overnight incubation at room temperature. Finally, it was centrifuged (10,000 g, 4 °C, 1 hour) and the pallet was redispersed in 1 mL of 1:1 H_2_O:HFIP (1,1,1,3,3,3-Hexafluoro-2-propanol, Merck, Cat. No. 18127) (v/v). The suspension was sonicated for 10 min and then centrifuged (5,000 g, 4 °C, 1 hour). The Aβ aggregate-mimics were then redispersed in 200 μL of 1:1 H_2_O:DMSO (v/v), and stored at −20 °C until use.

### Biotinylation of antibodies

Sulfo-NHS-LC-LC-Biotin (Thermo Scientific, Cat. No. 21338) was first dissolved in anhydrous DMSO at 10 mM. To the antibody (100 μg, at 1 mg/mL in PBS), 10 molar equivalents of sulfo-NHS-LC-LC-Biotin and 10 μL of 1M NaHCO_3_ were introduced. The reaction mixture was incubated at 37 °C for 1 hour, and then quenched by excess UltraPure™ 1M Tris, pH 8.0 (Invitrogen^TM^, Cat. No.15568025). Excess sulfo-NHS-LC-LC-Biotin was removed by an Zeba^TM^ Spin Desalting Column (40 kDa MWCO, ThermoFisher, Cat. No. 87766). The biotinylated antibody was then concentrated by an Amicon spin filter (50 kDa MWCO), and its concentration was determined by A_280_. The degree of labelling was determined using Pierce^TM^ Fluorescence Biotin Quantification Kit (ThermoFisher, Cat. No. 46610; **Supplementary Table 1**).

### Labelling antibodies with fluorescent dyes

Alexa Fluor™ 647 NHS Ester (Thermo Scientific, Cat. No. A20006) was first dissolved in anhydrous DMSO at 10 mM. To the antibody (100 μg, at 1 mg/mL in PBS), 10 molar equivalents of Alexa Fluor™ 647 NHS Ester and 10 μL of 1M NaHCO_3_ were introduced. The reaction mixture was incubated at 37 °C for 1 hour, and then quenched by excess UltraPure™ 1M Tris, pH 8.0 (Invitrogen^TM^, Cat. No.15568025). Excess dyes were removed by an Zeba^TM^ Spin Desalting Column (40 kDa MWCO, ThermoFisher, Cat. No. 87766). The dye-labelled antibody was then concentrated by an Amicon spin filter (50 kDa MWCO). The concentration and degree of labelling were determined by A_280_ and A_650_ (**Supplementary Table 1**).

### Conjugating antibodies to SiMoA beads

The SiMoA beads were functionalised with antibodies according to manufacturer’s instructions. Briefly, 4.2 × 10^8^ SiMoA 488-dyed singleplex beads (Quanterix, Cat. No. 104006, Lot No. 231817) were washed three times with 0.01 M NaOH, followed by three times with H_2_O. The beads were then resuspended in 280 μL of 25 mM cold MES buffer (pH 5). Meanwhile, EDC (ThermoFisher, Cat. No. A35391) and sulfo-NHS (ThermoFisher, Cat. No. A39269) were freshly dissolved in 25 mM cold MES buffer (pH 5) at 10 mg/mL and 40 mg/mL, respectively. The washed beads were subsequently activated by mixing 4.5 μL of EDC and 15 μL of sulfo-NHS. The mixture was incubated on a HuLa mixer at 4 °C for 30 minutes. Meanwhile, ∼100 μg of antibody was buffer-exchanged to 25 mM cold MES buffer (pH 5) using an Amicon filter (50 kDa MWCO). The activated beads were then washed once with 25 mM cold MES buffer. Next, 45 μg of buffer-exchanged antibody was introduced to the activated beads. The mixture was incubated on a HuLa mixer at room temperature for 30 minutes. The conjugated beads were then washed twice with the provided bead wash buffer (in the Homebrew 2.0 Development Kit, Quanterix, Cat. No. 101354) and blocked by the provided bead blocking buffer (in the Homebrew 2.0 Development Kit) on a HuLa mixer at room temperature for 40 min. Finally, the blocked beads were washed once with the bead wash buffer and then once with the provided bead diluent (in the Homebrew 2.0 Development Kit). The beads were then dispersed in 300 μL of bead diluent and stored at 4 °C until use. The coating efficiency of the beads, which suggests >99% of the antibodies were conjugated to the beads, was calculated by A_280_ of the supernatants before the 40-min blocking.

### Preparation of human post-mortem AD brain homogenate samples

The post-mortem human brain samples with no known history of neurological or neuropsychiatric symptoms other than AD, were acquired from the King’s College (with the approval of the London—Bloomsbury Research Ethics Committee; 16/LO/0508) and Edinburgh (REC 21/ES/0087) Brain Banks. The brain samples have been voluntarily donated without any compensation. The MTG segments were dissected and flash-frozen to be stored in −80℃ freezers until processing.

The samples were homogenised using a previously published method^2^. In brief, approximately 400 mg of tissue samples were placed into 2 ml tubes prefilled with 1 mm zirconium beads (Scientific Labs, Cat. SLS1414). 800 µl of homogenising buffer (10 mM Tris-HCl, 0.8 M NaCl, 1 mM EGTA, 10% sucrose, 0.1% sarkosyl, Pefabloc® SC protease inhibitor and PhosSTOP™ phosphatase inhibitor tablet; pH ∼7.4) was added and the samples were mechanically homogenised on an electronic tissue homogeniser (VelociRuptor V2 Microtube Homogeniser, Scientific Labs, Cat. SLS1401), at 5 meters/sec for 2 cycles of 15 seconds, with a 10 second gap in between, followed by centrifugation at 21,000 g for 20 minutes at 4℃. Then the supernatant was collected and stored in a LoBind Eppendorf at 4℃, while the pellet was homogenised again by adding an additional 800 µl of homogenising buffer and repeating the steps above. The supernatant from this step was mixed with the one from the previous step, aliquoted and stored in a −80℃ freezer.

### SiMoA sample preparation

Early oligomers, late oligomers, and fibrils made from *in-vitro* Aβ aggregates were diluted with the Homebrew sample/detector diluent provided in the Homebrew 2.0 Development Kit (Quanterix, Cat. No. 101354) to give the following concentrations based on monomeric equivalents: 9.0 nM, 2.3 nM, 562.5 pM, 140.6 pM, 35.2 pM, 8.8 pM, and 2.2 pM. Human-brain homogenate samples were diluted in 5:95 ratio with Lysate diluent A, as provided in the SiMoA lysate diluent kit (Quanterix, Cat. No. 102907).

### SiMoA protocol

SiMoA beads coated with Lecanemab, Aducanumab or Gantenerumab were first washed three times with the provided bead diluent in the Homebrew 2.0 Development Kit (Quanterix, Cat. No. 101354). Meanwhile, 100 µL of samples (see previous section) were loaded to each well on a conical bottom microplate (Quanterix). Then, 25 µL of the washed beads were introduced to each well to give the final-bead concentration as 2 × 10^7^ beads/mL. The samples and the beads were incubated on the plate shaker at 30 °C at 800 rpm for 1 hour. The plate was then washed by the SiMoA washer with the provided buffers. Afterwards, the biotinylated detector antibodies, i.e. biotinylated Lecanemab, Aducanumab or Gantenerumab, were then diluted to 37.5 ng/mL with the provided Homebrew sample/detector diluent and 100 µL of the diluted detectors were introduced to each well. The mixture was then incubated on the plate shaker at 30 °C with 800 rpm for 10 min. Similarly, the plate was then washed by the SiMoA washer. Meanwhile, the provided SBG concentrate was diluted to 25 pM with SBG diluent. To each well, 100 µL of the diluted SBG was introduced and the mixture was incubated on the plate shaker at 25 °C at 800 rpm for 10 min. Finally, the plate was washed by the SiMoA washer and loaded onto the SR-X™ Biomarker Detection System (Quanterix) together with the SiMoA discs, tips, and an equilibrated, well-shaken and opened RGP bottle.

### Immuno-blocking of human brain-derived Aβ aggregates

The pooled Braak Stage VI brain homogenate samples were first incubated with different concentrations of unlabelled Lecanemab, Aducanumab or Gantenerumab (0 nM, 2 nM, 20 nM, 200 nM and 2000 nM) in PBS on a HuLa mixer at room temperature for 1 hour. The samples were then diluted in 5:95 ratio with Lysate diluent A, as provided in the SiMoA lysate diluent kit.

### Single-molecule pull-down (SiMPull) coverslip preparation

Surface of the glass coverslips (VWR, Cat. No. MENZBC026076AC40) were passivated as previously described^3–5^. In brief, the coverslips were first cleaned by sonication (Ultrasonic cleaner USC100T, VWR) with a series of solvents (10 min in each of 18.2 MΩ cm water, acetone (Thermo Fisher, Cat. No. 10442631), and methanol (Thermo Fisher, Cat. No. 10675112)). The coverslips were then etched by sonication with 1 M KOH for 20 min. After that, they were rinsed with methanol, 18.2-MΩ·cm water, and methanol before dried with a stream of nitrogen. They were then cleaned with argon plasma (PDC-002, Harrick Plasma) for 15 min. The surfaces were then silanised with 3-aminopropyl triethoxysilane (Fisher Scientific UK, cat. no. 10677502), acetic acid (Merck, Cat. No. 45726) and methanol in the ratio of 3:5:100 in the sonicator for 60-s sonication with 10-min rest for two cycles. The coverslips were then rinsed with methanol, 18.2-MΩ·cm water, and methanol before dried with a stream of nitrogen. A 50-well PDMS gasket (Merck, Cat. No. GBL103250) was then affixed each coverslip. Each well was passivated by firstly introducing 9 μL of a freshly prepared 100:1 aqueous mixture of methoxy-PEG-Succinimidyl Valerate (110 mg/mL, Mw ∼5,000; Laysan Bio Inc., Cat. No. MPEG-SVA-5000) and biotin-PEG-Succinimidyl Valerate (100 mg/mL, Mw ∼5,000; Laysan Bio Inc., Cat. No. Biotin-PEG-SVA-5000) followed by adding 1 μL of 1 M NaHCO3 (pH 8.5). After overnight incubation in a humid chamber at room temperature, the coverslips were rinsed with 18.2-MΩ·cm water and dried with a stream of nitrogen. Each well was then further passivated by adding 9 μL of a freshly prepared aqueous solution of methyl-PEG4-NHS-Ester (10 mg/mL; Thermo Fisher, Cat. No. 22341), followed by adding 1 μL of 1 M NaHCO3 (pH 8.5). The coverslips were again incubated overnight in a humid chamber at room temperature. They were then washed with 18.2-MΩ·cm water and dried with a stream of nitrogen. Finally, they were stored in a desiccator at −20°C until use.

### SiMPull protocol

1× PBS containing 0.05% Tween 20 (Fisher Scientific, Cat. No. BP337-100; PBST) and 1× PBS containing 1% Tween 20 were freshly filtered by 0.02-μm filter (VWR, Cat. No. 516-1501) before use. *d*STORM buffer, which contains glucose oxidase (Merck, Cat. no. G2133), catalase (Merck, Cat. no. C40), cysteamine (Merck, Cat. no. 30070), and glucose in 50 mM Tris supplemented with 10 mM NaCl at pH 8. The glucose oxidase and catalase were first freshly dissolved in GOD buffer (24 mM PIPES, 4 mM MgCl_2_, 2 mM EGTA, pH 6.8) to give 40 mg/mL and 5 mg/mL stock solution, respectively. Meanwhile, a 25% glucose solution (in 18.2-MΩ·cm water) and 1M cysteamine solution (in 0.36 M HCl) were freshly prepared. The final working imaging buffer for *d*STORM contains 0.5 mg/mL glucose oxidase, 40 μg/mL catalase, 50 mM cysteamine, and 10% glucose in 50 mM Tris supplemented with 10 mM NaCl at pH 8. This solution was prepared freshly and immediately before imaging.

Unless otherwise specified, the wells were washed by a three-step protocol: twice with 10 μL PBST and once with 10 μL PBS containing 1% Tween 20. To each well on the biotinylated surface, 10 μL of NeutrAvidin (0.2 mg/mL in PBST, Thermo Fisher, Cat. No. 31000) was introduced and incubated for 10 min. Each well was then rinsed by the three-step protocol. Biotinylated antibodies, i.e., biotinylated Lecanemab, Aducanumab or Gantenerumab, were then diluted to 10 nM in 1× PBS supplemented with 0.1 mg/mL BSA (Thermo Fisher, Cat. No. B14) and 10 μL was added to each well for 10 min. Each well was then rinsed by the three-step protocol. Meanwhile, an aliquot of human-derived aggregates from the brain tissue was then diluted 10-fold (v/v) with 1× PBS or an aliquot of *in vitro* Aβ aggregates was diluted to 1 μM (monomeric equivalent) with 1× PBS or control sample, 1 mg/mL BSA in 1× PBS, was prepared. Next, 10 μL of diluted aggregates was added to each well for 90-min incubation. Each well was then rinsed by the three-step protocol. BSA was then diluted to 1 mg/mL in 1× PBS and 10 μL was added to each well for 30 min. Each well was then rinsed by the three-step protocol. Fluorescently labelled antibodies, i.e., AF647-labelled Lecanemab, Aducanumab or Gantenerumab, were then diluted to 1 nM in SmartBlock (CANDOR Bioscience GmbH, Cat. No. 113-125) and 10 μL was added to each well for 30 min. Each well was then rinsed by the three-step protocol. The solution in the well was then withdrawn and another two PDMS gaskets were stacked on and aligned to the PDMS gasket already on the coverslip. Each well was then first filled with 10 μL of 1× PBS for diffraction-limited imaging and subsequently replaced by 10 μL of *d*STORM buffer for super-resolution imaging. Finally, a cleaned glass coverslip was layered on top of the PDMS gaskets before super-resolution imaging.

### Microscope setup

Four lasers operating at 405 nm (LDM-405-350-C, Lasertack GmbH), 488 nm (Toptica iBeam smart, Toptica), 561 nm (Cobolt Jive, HÜBNER GmbH & Co KG) and 638 nm (Cobolt 06-MLD-638, HÜBNER GmbH & Co KG) were coupled to the optical axis of a 1.49 numerical aperture (NA) 100× CFI Apo TIRF objective (MRD01991, Nikon) mounted on an inverted Ti-E Eclipse microscope (Nikon, Japan). The lasers’ powers were controlled by their corresponding software or attenuated by neutral density filters. The laser beams were then passed through the aligning mirrors and were combined by their corresponding dichroic mirror (for 405 nm: FF458-Di02-25×36, Semrock; for 488 nm: FF552-Di02-25×36, Semrock; for 561 nm: FF605-Di02-25×36, Semrock) before being focused by an aspheric lens (C220TMDA, Thorlabs) to the square-core optical fibre (05806-1 Rev. A, CeramOptec). Launching was optimised using a free space fibre launch system (KT120/M, Thorlabs) and speckles from the fibre were removed using a vibration motor (304-111, Precision Microdrives Ltd.) mounted on a custom 3D printed mount^6^. The combined laser beam coming out from the optical fibre was then collimated (C40FC-A, Thorlabs) and cleaned up by a quad-band excitation filter (FF01-390/482/563/640-25×36). To mechanically decouple the collimator from the microscope body, one thick (OR26X2V175, Hooper Ltd.) Viton O ring was inserted between the external thread of the collimator and the internal thread of the mating optomechanics to minimise mechanical vibrations. The cleaned and collimated beam was then passed through the back port of the microscope and landed on an achromatic doublet lens (AC254-125-A-ML, Thorlabs). The excitation beam was then reflected by a penta-band dichroic beam splitter (R405/488/561/635/800-T1-25×36, Semrock) and focused on the sample by the objective. The size of the excitation beam was modified by adjusting the distance between the collimator (C40FC-A, Thorlabs) and back focal plane. Fluorescence from the sample was collected by the objective and passed through a quad-band emission filter (FF01-446/523/600/677-25×36, Semrock) and their corresponding appropriate filters (for both 405- and 488-nm induced fluorescence: BLP01-488R-25×36, Semrock and FF01-520/44-25×36, Semrock; for 561-nm induced fluorescence: LP02-568RS-25×36, Semrock and FF01-587/35-25×36, Semrock; for 638-nm induced fluorescence BLP01-635R-25×36, Semrock) mounted on a high-speed filter wheel (HF110A, Prior Scientific) before being recorded on an EMCCD camera (Evolve 512, Photometrics) operating in frame transfer mode (EM Gain of 3.1 electrons/ADU and 250 ADU/photon). Each pixel corresponded to a length of 101.2 nm on the recorded image. The microscope was also fitted with a perfect focus system (PFS) which auto-corrects the z-stage drift during a prolonged period of imaging. To remove the stray infrared laser beam from the PFS, a short-pass filter (FESH0750, Thorlabs) was mounted on the entrance port of the EMCCD camera.

### Image acquisition and analysis

Slides were fixed on a microscope stage and coupled to an objective using refractive index-matched low-autofluorescence immersion oil (refractive index n = 1.518, Olympus, UK). Images were taken in a grid using an automation script (μManager v1.4.22). Exposure times were set at 50 ms and 33 ms for diffraction-limited and super-resolution imaging, respectively. For diffraction-limited and super-resolution imaging, 50 frames and 5,000 frames were acquired, respectively. The power densities for diffraction-limited and super-resolution imaging were 0.11 kW/cm^2^ and 1.8 kW/cm^2^, respectively. The power density for the 405-nm activation laser for super-resolution imaging was 800 W/cm^2^. All data were analysed by the Aggregate Characterization Toolkit (ACT)^7^. For diffraction-limited images, the number and brightness of spots were determined by ComDet with the threshold as ‘3’ and estimated size as ‘5’. The number of spots was then subsequently filtered by the threshold set by the average IntPerArea determined by the data from control samples. For super-resolution images, all data were reconstructed with the scale set as ‘8’. Cross-correlation was applied for drift correction. Cluster analyses were conducted by DBSCAN, with EPS set as ‘75 nm’ and Min sample set as ‘2’.

### Statistical analyses

The number of replicates for each experiment and the statistical values are provided in the corresponding figure captions. The R Project Statistical Computing version 4.4.1 (2024-06-14) -- “Race for Your Life” was used for all statistical analyses, the graphs were generated in GraphPad Prism 7.0a for Mac OS X, and the cartoon figures were created with BioRender.com. ANOVA with Type 2 sums of squares were used to determine the differences between the groups. Data collected during this experiment will be made available upon reasonable request from the corresponding author. No novel code or software were generated to collect or analyse the data.

## Main text references

1. Gustavsson, A. et al. Global estimates on the number of persons across the Alzheimer’s disease continuum. Alzheimers Dement 19, 658–670 (2023).

2. Selkoe, D. J. Amyloid protein and Alzheimer’s disease. Scientific American 265, 68–71, 74–76, 78 (1991).

3. Hardy, J. & Allsop, D. Amyloid deposition as the central event in the aetiology of Alzheimer’s disease. Trends in pharmacological sciences 12, 383–388 (1991).

4. Arndt, J. W. et al. Structural and kinetic basis for the selectivity of aducanumab for aggregated forms of amyloid-β. Sci Rep 8, 6412 (2018).

5. Swanson, C. J. et al. A randomized, double-blind, phase 2b proof-of-concept clinical trial in early Alzheimer’s disease with lecanemab, an anti-Aβ protofibril antibody. Alzheimers Res Ther 13, 80 (2021).

6. Bohrmann, B. et al. Gantenerumab: a novel human anti-Aβ antibody demonstrates sustained cerebral amyloid-β binding and elicits cell-mediated removal of human amyloid-β. J Alzheimers Dis 28, 49–69 (2012).

7. Rashad, A. et al. Donanemab for Alzheimer’s Disease: A Systematic Review of Clinical Trials. Healthcare (Basel) 11, 32 (2022).

8. Knopman, D. S. Lecanemab reduces brain amyloid-β and delays cognitive worsening. Cell Rep Med 4, 100982 (2023).

9. Söderberg, L. et al. Lecanemab, Aducanumab, and Gantenerumab — Binding Profiles to Different Forms of Amyloid-Beta Might Explain Efficacy and Side Effects in Clinical Trials for Alzheimer’s Disease. Neurotherapeutics 20, 195–206 (2023).

10. Sims, J. R. et al. Donanemab in Early Symptomatic Alzheimer Disease: The TRAILBLAZER-ALZ 2 Randomized Clinical Trial. JAMA 330, 512–527 (2023).

11. Danial, J. S. H. & Klenerman, D. Single molecule imaging of protein aggregation in Dementia: Methods, insights and prospects. Neurobiology of Disease 153, 105327 (2021).

12. Fertan, E. et al. Cerebral organoids with chromosome 21 trisomy secrete Alzheimer’s disease-related soluble aggregates detectable by single-molecule-fluorescence and super-resolution microscopy. Mol Psychiatry (2023) doi:10.1038/s41380-023-02333-3.

13. Sideris, D. I. et al. Soluble amyloid beta-containing aggregates are present throughout the brain at early stages of Alzheimer’s disease. Brain communications 3, (2021).

14. De, S. et al. Different soluble aggregates of Aβ42 can give rise to cellular toxicity through different mechanisms. Nature Communications 2019 10:1 10, 1–11 (2019).

15. McDade, E. et al. Lecanemab in patients with early Alzheimer’s disease: detailed results on biomarker, cognitive, and clinical effects from the randomized and open-label extension of the phase 2 proof-of-concept study. Alzheimers Res Ther 14, 191 (2022).

16. Whitehouse, P. J. & Saini, V. Making the Case for the Accelerated Withdrawal of Aducanumab. J Alzheimers Dis 87, 999–1001 (2022).

17. Bateman, R. J. et al. Two Phase 3 Trials of Gantenerumab in Early Alzheimer’s Disease. N Engl J Med 389, 1862–1876 (2023).

18. Julku, U. et al. Brain pharmacokinetics of mono- and bispecific amyloid-β antibodies in wild-type and Alzheimer’s disease mice measured by high cut-off microdialysis. Fluids and Barriers of the CNS 19, 99 (2022).

## Online methods’ references

1. Hülsemann, M. et al. Biofunctionalized Silica Nanoparticles: Standards in Amyloid-β Oligomer-Based Diagnosis of Alzheimer’s Disease. J Alzheimers Dis 54, 79–88 (2016).

2. Böken, D. et al. Single-Molecule Characterization and Super-Resolution Imaging of Alzheimer’s Disease-Relevant Tau Aggregates in Human Samples. Angew Chem Int Ed Engl e202317756 (2024) doi:10.1002/anie.202317756.

3. Fertan, E. et al. Cerebral organoids with chromosome 21 trisomy secrete Alzheimer’s disease-related soluble aggregates detectable by single-molecule-fluorescence and super-resolution microscopy. Mol Psychiatry (2023) doi:10.1038/s41380-023-02333-3.

4. Antonio, M. D. et al. Single-molecule visualization of DNA G-quadruplex formation in live cells. Nature chemistry 12, 832–837 (2020).

5. Jain, A. et al. Probing cellular protein complexes using single-molecule pull-down. Nature 473, 484–488 (2011).

6. Lam, J. Y. L., et al. An economic, square-shaped flat-field illumination module for TIRF-based super-resolution microscopy. Biophys Rep (N Y) 2, None (2022).

7. Xia, Z. et al. A computational suite for the structural and functional characterization of amyloid aggregates. Cell Reports Methods 3, 100499 (2023).

