## Supplemental Table 1 for "Single-molecule characterisation of soluble beta-amyloid aggregate binding by Aducanumab, Lecanemab, Gantenerumab, and Donanemab"

**Supplemental Table 1. The labelling efficiency of the antibodies.**

| MAB | No. of biotins / MAb | No. of dyes (Alexa Fluor 647) / MAb |
|---|---|---|
| Aducanumab | 1.3 | 6.0 |
| Lecanemab | 1.0 | 4.7 |
| Gantenerumab | 1.2 | 5.5 |
| Donanemab | 2.2 | 4.1 |
